# A xylose-inducible expression system and a CRISPRi-plasmid for targeted knock-down of gene expression in *Clostridioides difficile*

**DOI:** 10.1101/476796

**Authors:** Ute Müh, Anthony G. Pannullo, David S. Weiss, Craig D. Ellermeier

**Author notes:** Corresponding Author: Craig D. Ellermeier,; David S. Weiss.

## Abstract

Here we introduce plasmids for xylose-regulated expression and repression of genes in *Clostridioides difficile*. The xylose-inducible expression vector allows for ~100-fold induction of an *mCherryOpt* reporter gene. Induction is titratable and uniform from cell-to-cell. The gene repression plasmid is a CRISPR-interference (CRISPRi) system based on a nuclease-defective, codon-optimized allele of the *Streptococcus pyogenes* Cas9 protein (*dCas9*) that is targeted to a gene of interest by a constitutively-expressed single guide RNA (sgRNA). Expression of *dCas9* is induced by xylose, allowing investigators to control the timing and extent of gene-silencing, as demonstrated here by dose-dependent repression of a chromosomal gene for a red fluorescent protein (maximum repression ~100-fold). To validate the utility of CRISPRi for deciphering gene function in *C. difficile*, we knocked-down expression of three genes involved in biogenesis of the cell envelope: the cell division gene *ftsZ*, the S-layer protein gene *slpA* and the peptidoglycan synthase gene *pbp-0712*. CRISPRi confirmed known or expected phenotypes associated with loss of FtsZ and SlpA, and revealed that the previously uncharacterized peptidoglycan synthase PBP-0712 is needed for proper elongation, cell division and protection against lysis.

**Importance:** *Clostridioides difficile* has become the leading cause of hospital-acquired diarrhea in developed countries. A better understanding of the basic biology of this devastating pathogen might lead to novel approaches for preventing or treating *C. difficile* infections. Here we introduce new plasmid vectors that allow for titratable induction (P_*xyl*_) or knockdown (CRISPRi) of gene expression. The CRISPRi plasmid allows for easy depletion of target proteins in *C. difficile*. Besides bypassing the lengthy process of mutant construction, CRISPRi can be used to study the function of essential genes, which are particularly important targets for antibiotic development.

## Introduction

*Clostridioides* (formerly *Clostridium*) *difficile* is a strictly anaerobic, Gram-positive opportunistic pathogen that has become the leading cause of healthcare-associated diarrhea (1). The symptoms are caused by toxins that damage the intestinal epithelium and can range in severity from mild diarrhea to life threatening conditions such as pseudomembranous colitis and toxic megacolon (2–4). A recent study estimated that *C. difficile* causes close to ~500,000 infections and contributes to ~25,000 deaths per year in the United States (1, 5). The Centers for Disease control has classified *C. difficile* as an “urgent threat” to human health (6).

Genetic tools to study the physiology of *C. difficile* have become increasingly available, nevertheless the repertoire remains limited. Investigators wishing to manipulate gene expression have two options, a tetracycline-inducible promoter, P_*tet*_ (7), and a nisin-inducible promoter P_*cpr*_ (8, 9). Although both systems are useful, the inducers, anhydrotetracycline and nisin, can inhibit growth (8, 10). This can make it difficult to deconvolute the effects of the inducer from the effects of altered gene expression. Investigators wishing to construct targeted knock-outs in *C. difficile* have several options (11–13), but none of these can be applied to determine the function of essential genes and even in the case of non-essential genes the effort required to construct and validate mutants remains a significant bottleneck.

Here we describe two new plasmids that address some of these limitations. The first is a xylose-inducible expression vector based on the native P_*xyl*_ promoter of *C. difficile* (pAP114, Fig. 1A). Similar xylose-inducible expression vectors have been developed for a number of bacteria (14–16), including one for *Clostridium perfringens* that is based on the *C. difficile* xylose-regulatory system (17). The xylose-inducible vector allows for uniform, tunable expression of an *mCherryOpt* reporter gene, with a maximum induction ratio of >100-fold.

**Fig. 1.**
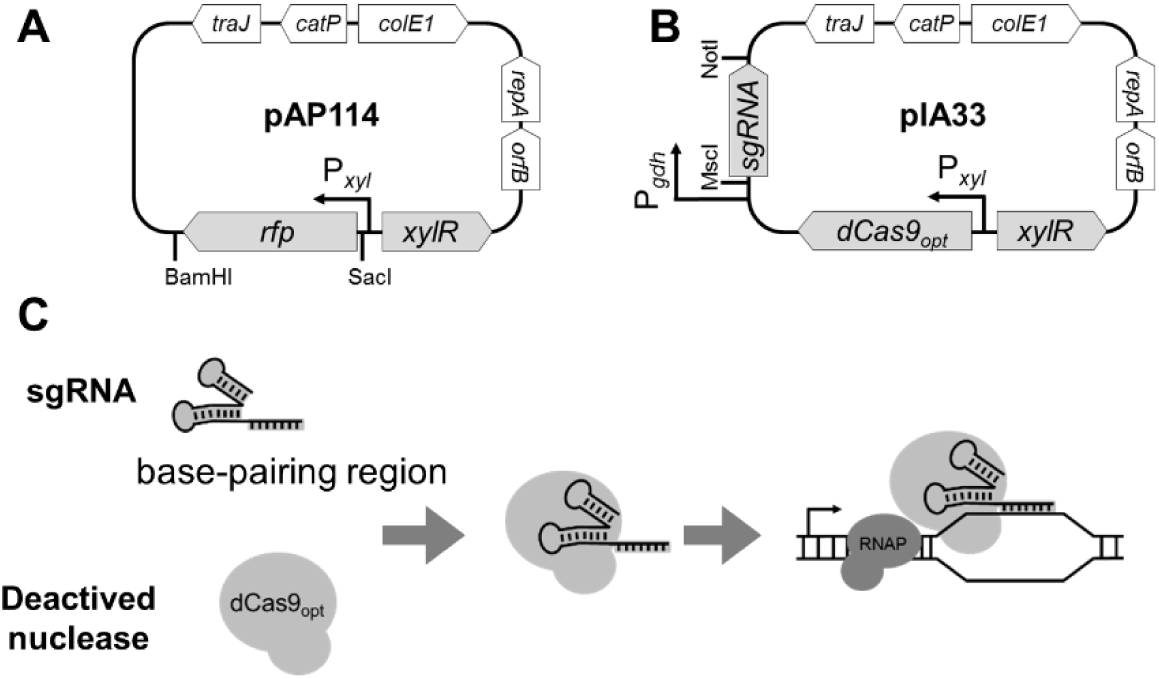
Genetic maps and CRISPRi principle. (A) pAP114 for xylose inducible expression. Expression of *rfp* is under P_*xyl*_ control. The gene can be replaced using the SacI and BamHI restriction sites. (B) pIA33 for CRISPRi. Single guide RNA (sgRNA) is under P_*gdh*_ control and *dCas9* is under P_*xyl*_ control. The *sgRNA* sequence can be replaced using the MscI and NotI restriction sites. (C) CRISPRi principle. The deactivated nuclease binds sgRNA and is targeted to the gene of interest, but is unable to cleave the DNA. Instead dCas9 serves as a transcriptional roadblock. Features depicted: *rfp* encodes a red fluorescent protein variant called mCherryOpt that is codon optimized for expression in low-GC bacteria; *dCas9* encodes the deactivated CRISPR nuclease codon-optimized for expression in low-GC bacteria; *orfB* and *repA*, the replication region; *colE1*, replication region of the *E. coli plasmid* pBR322 modified for higher copy number; *catP*, the chloramphenicol acetyltransferase gene from *Clostridium perfringens*, conferring resistance to thiamphenicol in *C. difficile* or chloramphenicol in *E. coli*; *traJ*, encodes a conjugation transfer protein from plasmid RP4. Modified from (23).

The second new plasmid is a CRISPR interference (CRISPRi) vector that allows investigators to quickly assess the function of target genes, including essential genes, by blocking their expression (pIA33, Fig. 1B). CRISPRi has been used to explore gene function in a variety of bacteria (e.g., (18–22)), we modeled our plasmid after a CRISPRi tool developed for *Bacillus subtilis* (23). CRISPRi uses a single guide RNA (sgRNA) to deliver a nuclease-deactivated mutant of Cas9 (dCas9) to a gene of interest, thereby creating a roadblock that prevents transcription by RNA polymerase (Fig. 1C) (20).

Targeting repression to a new gene is simply a matter of replacing the sgRNA. Because *dCas9* is expressed from P_*xyl*_, investigators can use xylose to control the timing and extent of gene silencing. To validate the utility of the CRISPRi vector, we show it can achieve tunable repression of a chromosomal gene for a red fluorescent protein (*rfp*) and document phenotypic defects caused by knocking down expression of three genes that are critical for biogenesis of the cell envelope: *ftsZ*, *slpA* and *cdr20291_0712,* which encodes a penicillin-binding protein referred to here as PBP-0712.

## Results

### Construction of a xylose-inducible expression system for *C. difficile*

To circumvent the deleterious effects of inducer toxicity inherent in the P_*tet*_ system (10), we constructed a xylose-inducible expression system. In *C. difficile*, the xylose gene cluster comprises the xylose repressor, *xylR* (*cdr20291_2900*), and the divergently-transcribed catabolic genes, *xylBA (24)*. As determined in a variety of Gram positive organisms, in the absence of xylose, XylR binds to the *xyl* operator (*xylO*) to repress transcription at P_*xyl*_ (24–26). To construct a xylose-inducible expression vector, we PCR-amplified a 1.4 kb *xylR*-P_*xyl*_ DNA fragment from the chromosome of *C. difficile* strain R20291. This fragment was cloned upstream of a codon-optimized gene for the red fluorescent protein mCherry to create the P_*xyl*_::*mCherryOpt* reporter plasmid named pAP114 (Fig. 1A).

R20291 harboring pAP114 exhibited a dose-dependent increase in red fluorescence when grown in media containing increasing xylose concentrations, achieving an induction range of >100-fold (Fig. 2A). Flow cytometry revealed the bacterial population was uniformly fluorescent over the full range of xylose concentrations (Fig. 2B). As expected, xylose did not inhibit growth even at the highest concentration tested, 3% (Fig. 2A, inset). Similar results were obtained when pAP114 was introduced into *C. difficile* strain 630Δ*erm* (Fig. S1).

**Fig. 2.**
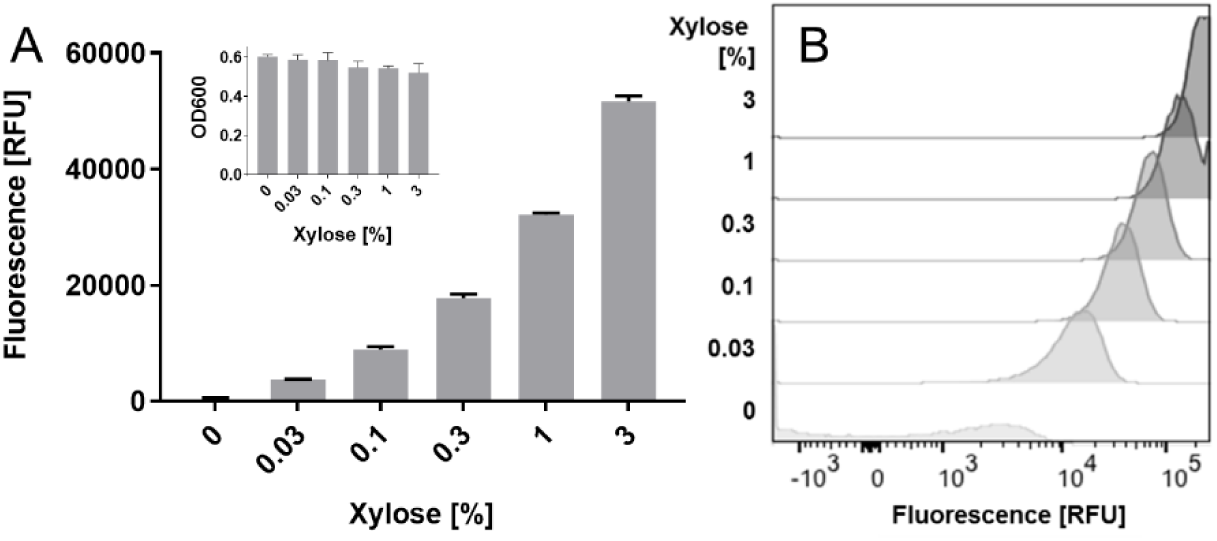
Tunable induction from P_*xyl*_. An overnight culture of R20291/pAP114 was diluted to a starting OD_600_ = 0.05 into TY Thi_10_ with the indicated concentration of xylose. Once cells had reached an OD_600_ = 0.5 (~5h) they were fixed and processed to allow RFP development. (A) A plate reader was used to measure relative fluorescence and OD_600_ of bulk samples. (B) Flow cytometry was used to measure fluorescence of individual cells. RFU is relative fluorescence units normalized to OD_600_. Data in (A) represent the mean and standard deviation of triplicate cultures. These results are representative of at least two independent experiments.

Expression of sugar utilization genes is often subject to carbon catabolite repression and in *C. difficile* is controlled by CcpA. Antunes *et al.* found that *xylA* expression was 3-fold lower in a *ccpA* mutant but was not subject to glucose repression (27). The *xylA* gene is located downstream of *xylB*. Thus, we sought to determine if expression of P*_xyl_* was subject to glucose repression. We found that xylose induction of P*_xyl_* decreased by less than 40% in the presence of glucose, confirming that glucose does not play a major role in regulation of this promoter (Fig. S2) (27).

### Construction of a CRISPRi system for silencing gene expression in *C. difficile*

Peters *et al.* used knock-down of a chromosomal *rfp* allele expressed from the constitutive P_*veg*_ promoter to establish a CRISPRi system for *B. subtilis* (23). We took advantage of their P_*veg*_::*rfp* construct and *rfp-*targeting sgRNA to establish an analogous CRISPRi system for *C. difficile*. (Note: this *rfp* allele is different from the *mCherryOpt rfp* allele used above to study xylose induction in pAP114.) Our first step was to construct a reporter strain by integrating P_*veg*_::*rfp* downstream of *pyrE* to create strain UM275 (*C. difficile* 630Δ*erm* P_*veg*_::*rfp*).

We then constructed a series of *E. coli-C. difficile* shuttle vectors that expressed *Streptococcus pyogenes dCas9* under control of P_*xyl*_ and the *rfp*-targeted sgRNA (*rfp*^sgRNA^) under control of various constitutive promoters: P_*veg*_, P_*sigA*_ and P_*gdh*_ (12, 23, 28). As a negative control we use a pseudo sgRNA with 20 nt that do not anneal next to a PAM site (P_*gdh*_::*neg*^sgRNA^). The plasmids were conjugated into the reporter strain and gene silencing was quantified by measuring red fluorescence with and without induction of *dCas9*. RFP signal was reduced by each of the three constructs that expressed the *rfp*-targeting sgRNA, but not by the negative control (Fig. S3). Inhibition correlated with the concentration of xylose (i.e. induction of *dCas9*) and was strongest when the sgRNA ^was cloned behind P^*veg* and P_*gdh*_. We chose to continue with P_*gdh*_ where maximal inhibition was about 80% at 1% xylose (Fig. 3A and S3).

**Fig. 3.**
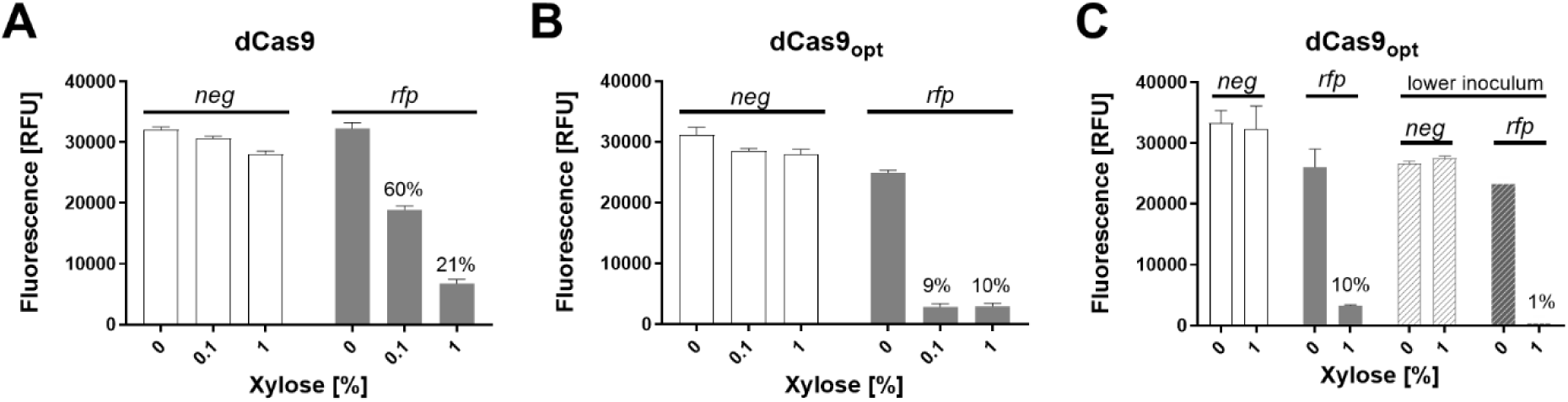
Suppression of *rfp* expression by CRISPRi. A set of CRISPRi plasmids was introduced into a *C. difficile* strain that expresses *rfp* constitutively. The CRISPRi plasmids express sgRNAs constitutively under P_*gdh*_ control and *dCas9* alleles under P_*xyl*_ control. Overnight cultures were diluted into TY Thi_10_ containing xylose as indicated. When cultures reached an OD_600_ = 0.5 (~5h), cells were fixed and processed to allow RFP development. In (A) cultures were inoculated to a starting OD_600_ = 0.05 and the CRISPRi plasmids expressed *dCas9* together with an *rfp*_sgRNA_ or a *neg_s_*^gRNA^ (negative control). (B) The *dCas9* gene was replaced with a codon-optimized *dCas9*. (C) The same CRISPRi plasmids as in (B) but cultures were inoculated to a starting OD_600_ = 0.05 or 0.01 (lower inoculum). Data are graphed as the mean and standard deviation of triplicate cultures and are representative of at least two independent experiments. The host strain was UM275. The plasmids were pAI25 (P_*gdh*_::*neg*^sgRNA^, P_*xyl*_::*dCas9*), pAI28 (P_*gdh*_::*rfp*^sgRNA^, P_*xyl*_::*dCas9*), pAI33 (P_*gdh*_::*rfp*^sgRNA^, P_*xyl*_::*dCas9-opt*), pAI34 (P_*gdh*_::*neg*^sgRNA^, P_*xyl*_::*dCas9-opt*).

We achieved further improvement by replacing the *dCas9* gene with one that had been codon-optimized for *C. difficile* (12) to create pIA33 (Fig. 1B). This final construct achieved 90% suppression of red fluorescence (Fig. 3B) when the reporter strain was grown for ~3 mass doublings (OD_600_ 0.05 to 0.5). The residual 10% RFP signal likely reflects carry-over from the inoculum, in which case the true repression would be greater than 90%. Consistent with this inference, reducing the inoculum lowered the residual fluorescence to ~1% (Fig. 3C).

### Applications of CRISPRi to study gene function in *C. difficile*

Once we optimized the CRISPRi plasmid against RFP, we tested its utility for studies of gene function by targeting three genes that play important roles in assembly of the cell envelope: *ftsZ*, *slpA*, and *cdr20291_0712* (*pbp-071*2). These genes were considered good candidates for CRISPRi because they are not co-transcribed with other genes (29, 30), minimizing concerns with polarity. Moreover, all three were identified as essential by saturation transposon mutagenesis (31), although a mutant lacking *slpA* was subsequently isolated by a different approach (32).

#### CRISPRi targeting *ftsZ*

FtsZ is a key division protein found in almost all bacteria (33, 34). Depletion of FtsZ in rod-shaped bacteria prevents division, leading to the formation of long filaments that eventually lyse. The appearance of highly filamentous cells provides a simple visual read-out that has been used to demonstrate successful inhibition of *ftsZ* expression by CRISPRi in *E. coli* (35, 36), *B. subtilis* (23), and *Pseudomonas aeruginosa* (22). In the coccus *Streptococcus pneumoniae*, CRISPRi knockdown of *ftsZ* expression causes swelling rather than filamentation (19).

We constructed three CRISPRi plasmids with different sgRNAs targeting *ftsZ* and conjugated these plasmids into strain R20291. All three constructs showed a strong growth-inhibition phenotype, with about 100 to 1000-fold loss in viability when serial dilutions of an overnight (stationary phase) culture were spotted onto TY Thi_10_ plates containing 1% xylose (Fig. 4A). Cells recovered from this plate were strikingly elongated, whereas the negative control exhibited normal morphology (Fig. S4). Liquid cultures of R20291 harboring one of the CRISPRi constructs were grown in the presence and absence of xylose and examined by microscopy after 4h of induction. Knockdown of *ftsZ* expression produced filamentous cells that lacked cell division septa (Fig. 4B). Collectively, these findings indicate successful silencing of *ftsZ* expression by CRISPRi.

**Fig. 4.**
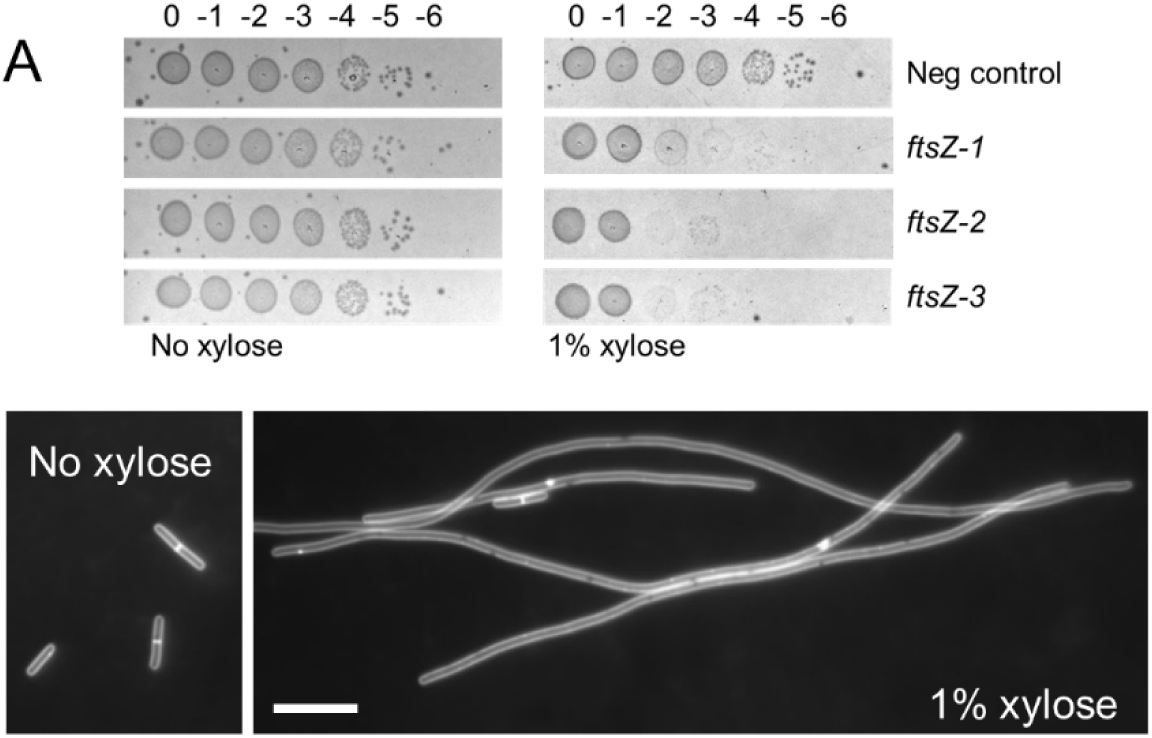
CRISPRi knockdown of *ftsZ*. (A) Viability assay. Cells of strain R20291 harboring CRISPRi plasmids were grown overnight in TY Thi_10_. Samples were serially diluted and 5 µl of each dilution were spotted onto TY Thi_10_ plates with or without 1% xylose to induce expression of *dCas9-opt*. Plates were photographed after incubation overnight. The negative control was pIA34 (*neg*^sgRNA^), while the plasmids that target *ftsZ* were pIA35 (*ftsZ-1*^sgRNA^), pIA36 (*ftsZ-2*^sgRNA^), and pIA37 (*ftsZ-3*^sgRNA^). (B) Cell morphology. An overnight culture of R20291/pAI35 (*ftsZ-1*^sgRNA^) was diluted to a starting OD_600_ = 0.05 in TY Thi_10_ with or without 1% xylose and grown for 4h. Cells were stained with the membrane dye FM4-64 to reveal whether division septa were present and photographed under fluorescence. Size bar = 10 µm. These results are representative of two independent experiments.

#### CRISPRi targeting *slpA*

The outermost layer of *C. difficile* is called the S-layer and consists of a two-dimensional paracrystalline array composed primarily of the <u>s</u>urface layer <u>p</u>rotein A, SlpA (37, 38). SlpA is post-translationally processed into a high molecular weight (HMW) and a low molecular weight (LMW) subunit that together form a tight heterodimeric complex which is incorporated into the S-layer (39, 40). It was initially thought that *slpA* might be essential since extensive transposon mutagenesis did not yield any insertions in *slpA* (31). However, growth in the presence of the SlpA-targeting, bactericidal agent Avidocin selected for strains that lacked the ability to produce SlpA (32). A *slpA* mutant strain obtained by this selection showed reduced sporulation, increased sensitivity to lysozyme and was attenuated in virulence (32). We set out to determine whether CRISPRi knockdown of *slpA* expression would confirm the phenotypes described for the *slpA*-mutant strain.

We constructed two CRISPRi plasmids with sgRNAs designed to prevent expression of *slpA*. R20291 harboring these plasmids was grown to stationary phase in the presence of 1% xylose. To assess the extent of SlpA depletion, cell wall proteins were extracted with low pH buffer, and samples were analyzed by SDS-PAGE. Based on the intensity of Coomassie staining, SlpA levels were reduced to less than 5% of wild-type (Fig. 5B). Because SlpA-depleted cells tended to clump, which might affect the efficiency of cell wall protein extraction, we further verified extensive depletion of SlpA in whole cell extracts (Fig. S5). Interestingly, the induced CRISPRi-*slpA* samples appear to lyse more readily during sample workup, as indicated by the higher levels of background bands.

**Fig. 5.**
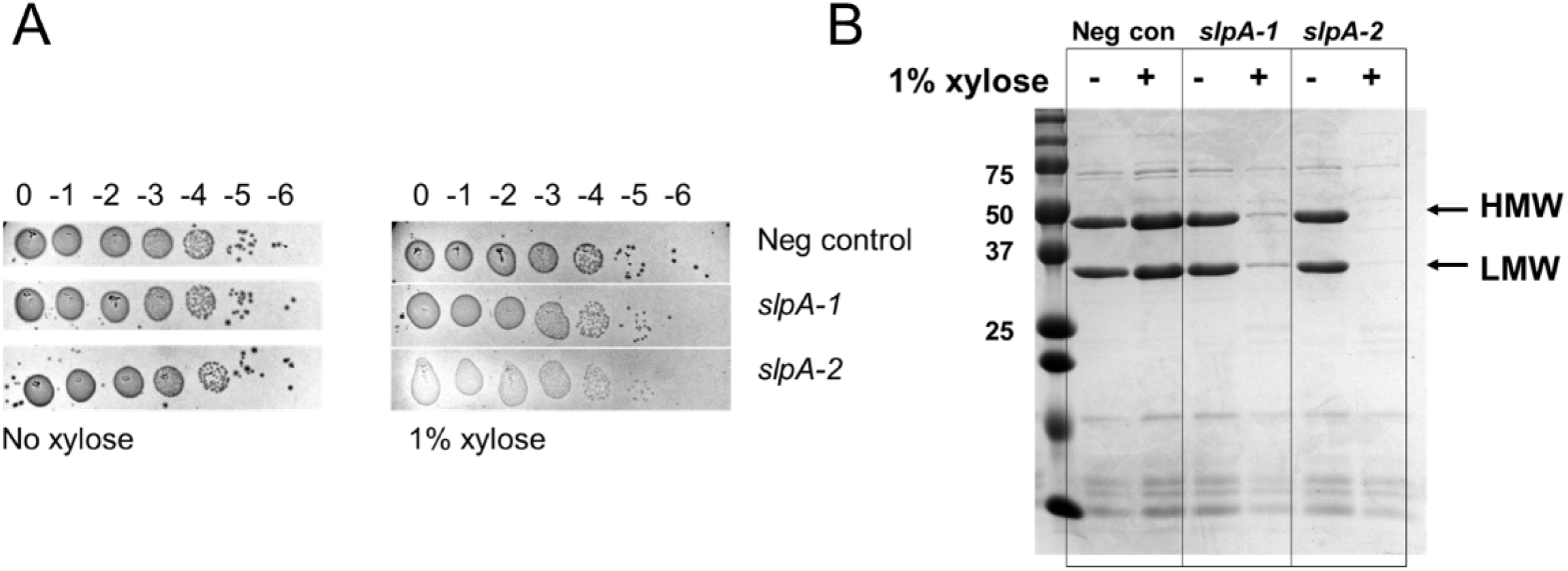
CRISPRi knockdown of *slpA*. (A) Viability assay as described in legend to Fig. 4. CRISPRi constructs used in this experiment were pIA34 (*neg*^sgRNA^; negative control), pIA38 (*slpA-1*^sgRNA^), or pIA39 (*slpA-2*^sgRNA^). Note: sgRNAs targeting *slpA* produced small, translucent colonies that did not photograph well and were difficult to score, but growth was consistently observed out to the 10^-5^ dilution in all cases. (B) Extent of SlpA depletion. The same strains were diluted 1:50 into TY Thi_10_ with or without 1% xylose. After growth for 8h cell wall proteins were extracted and analyzed by SDS-PAGE followed by Coomassie staining. Molecular mass markers in kilodaltons are indicated to the left. The high molecular weight band (HMW) and the low molecular weight band (LMW) of SlpA are marked by arrows. Results are representative of two experiments.

CRISPRi knockdown of *slpA* in R20291 did not affect plating efficiency, although the colonies that grew were more translucent than wild-type (Fig. 5A). SlpA-depleted cells exhibited normal rod morphology under phase contrast (Fig. S6). We found that knockdown of *slpA* expression led to increased lysozyme sensitivity in a growth curve at 0.2 mg/mL lysozyme (Fig. S7). Furthermore, the MIC for lysozyme was 8-16 mg/mL (Table 3) in a negative control strain or an uninduced CRISPRi-*slpA* strain, which decreased to 4 mg/mL when the CRISPRi-*slpA* construct was induced. Finally, CRISPRi silencing of *slpA* reduced sporulation on TY Thi_10_ plates about 1000-fold (Table 4). Our findings are fully consistent with those reported for a *slpA* null mutant obtained by selecting for resistance to Avidocin (32).

**TABLE 3.**
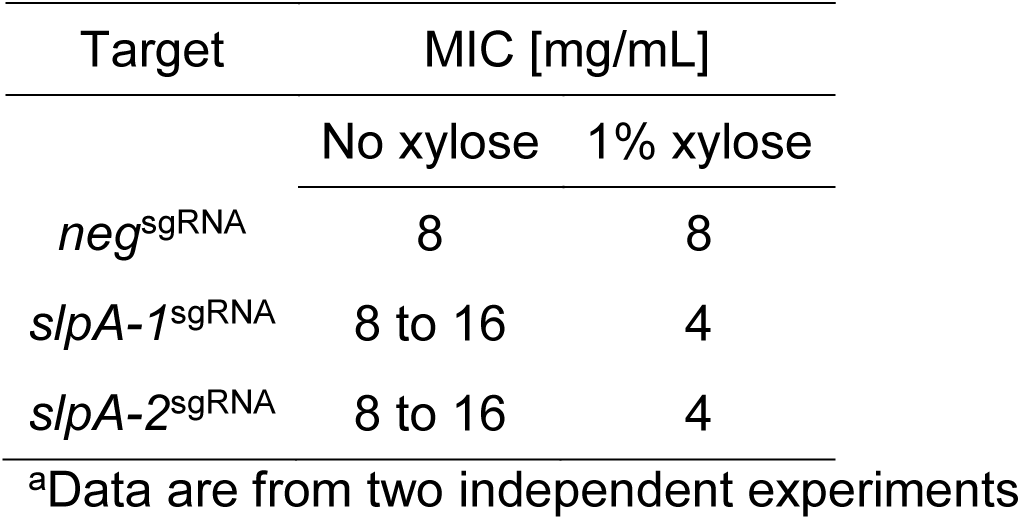
Effect of CRISPRi knockdown of *slpA* on MIC for lysozyme^a^

| Target | MIC [mg/mL] |  |
| --- | --- | --- |
|  | No xylose | 1% xylose |
| *neg*<sup>sgRNA</sup> | 8 | 8 |
| *slpA-1*<sup>sgRNA</sup> | 8 to 16 | 4 |
| *slpA-2*<sup>sgRNA</sup> | 8 to 16 | 4 |

<sup>a</sup>Data are from two independent experiments

**TABLE 4.** Effect of CRISPRi knockdown of *slpA* on sporulation^a^

| Target | Spores [%] |  | Fold reduction |
| --- | --- | --- | --- |
|  | No xylose | 1% xylose |  |
| *neg*<sup>sgRNA</sup> | 1.8 | 0.3 | 1 |
| *slpA-1*<sup>sgRNA</sup> | 0.5 | <0.002 | >250 |
| *slpA-2*<sup>sgRNA</sup> | 0.3 | <0.0005 | >600 |

<sup>a</sup>Data are the average of two independent experiments

### CRISPRi targeting *pbp-0712*

Penicillin-binding proteins (PBPs) are required to synthesize the peptidoglycan (PG) cell wall that surrounds most bacteria and protects them from lysis due to turgor pressure (41). Saturation transposon mutagenesis identified two PBPs as essential in *C. difficile* (31). One of these, *cdr20291_0985*, is not very amenable to analysis by CRISPRi because it is embedded in an operon with homologs of numerous genes known to be essential for division and elongation in other rod-shaped bacteria, including *B. subtilis* and *E. coli*. The other, coding for PBP-0712 *(cdr20291_0712)*, appears to be an isolated gene (29, 30). PBP-0712 is predicted to be a bifunctional (class A) PBP with both a glycosyltransferase domain for polymerization of glycan strands and a transpeptidase domain for crosslinking of adjacent stem peptides (42). Nothing is known about the specific roles of PBP-0712, such as whether it contributes primarily to elongation or division.

To determine the phenotype of PBP-0712 depletion, two CRISPRi plasmids with sgRNAs targeting *pbp-0712* were introduced into R20291 by conjugation. Exconjugates plated on TY Thi_10_ containing 1% xylose exhibited an approximate 10^6^-fold viability defect (Fig. 6A), confirming that *pbp-0712* is an essential gene. When inoculated to a starting OD_600_ = 0.05 and incubated for 4h, cultures grown without xylose reached an OD_600_ ~ 0.6 and the cells exhibited normal rod morphology (Fig 6B, inset). In contrast, cultures grown in the presence of 1% xylose reached an OD_600_ ~0.5 and exhibited a variety of morphological defects (Fig. 6B). Phase contrast microscopy revealed cell debris and lysed cells that looked like “ghosts,” as well as a mix of normal length and elongated rods, some of which were bent. Staining with the membrane dye FM4-64 revealed that a few of the filamentous cells had extensive regions lacking division septa, but most were chains of relatively short cells that appeared to have synthesized a division septum but not separated (a “chaining” phenotype). These morphological defects are reminiscent of those observed in a *C. difficile mld* mutant (43). Collectively, they implicate PBP-0712 in elongation, division and overall integrity of the PG sacculus.

**Fig. 6.**
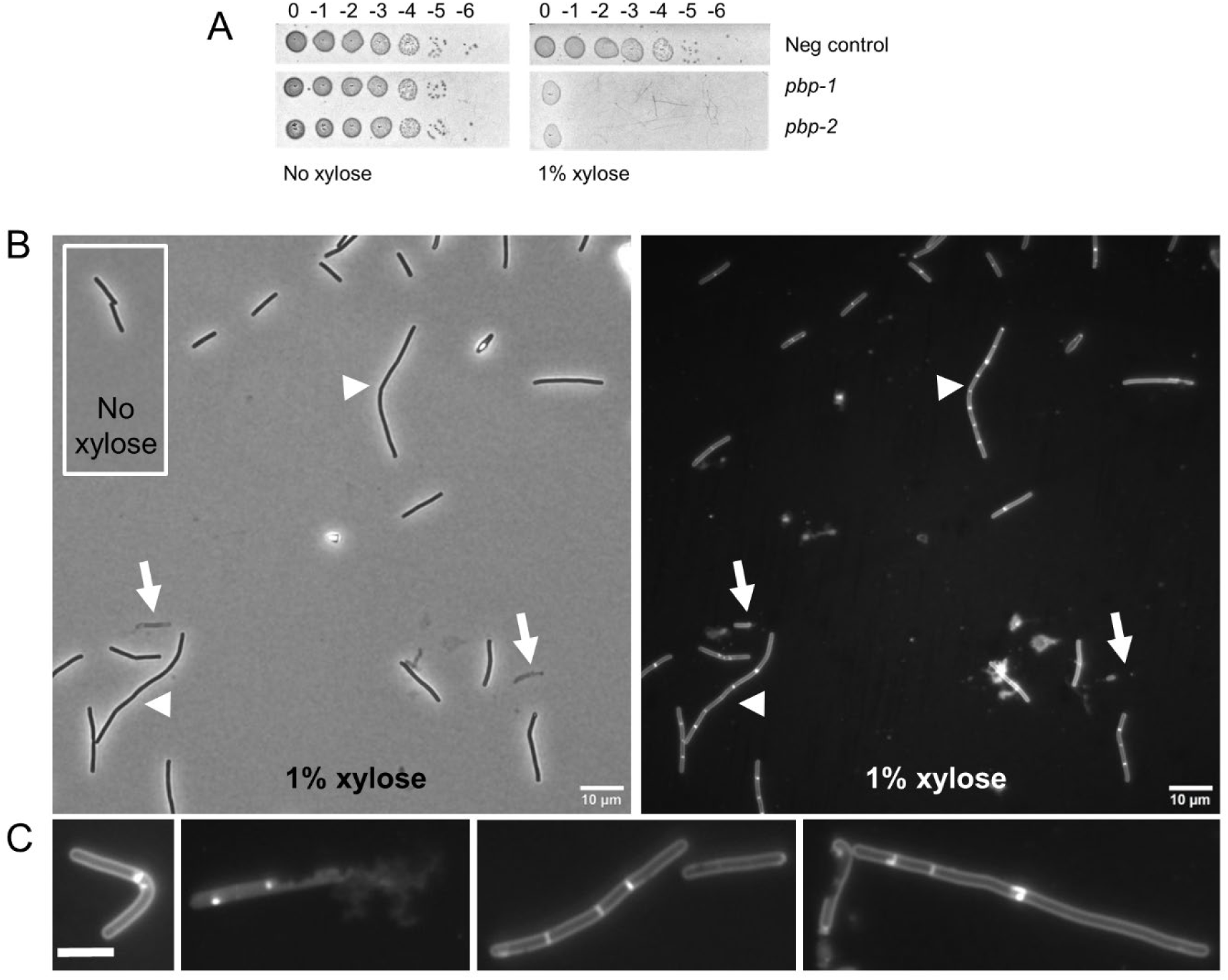
CRISPRi knockdown of *pbp-0712*. (A) Viability assay as described in legend to Fig. 4. CRISPRi constructs used in this experiment were pIA34 (*neg*^sgRNA^; negative control), pIA40 (*pbp-0712-1*^sgRNA^), or pIA41 (*pbp-0712-2*^sgRNA^). (B) Cell morphology. An overnight culture of R20291/pIA40 (*pbp-0712-1*^sgRNA^) grown in TY Thi_10_ was inoculated to a starting OD_600_ = 0.05 in TY Thi_10_ with or without 1% xylose. After growth for 6h cells were stained with the membrane dye FM4-64 and photographed under phase contrast (left) or fluorescence (right). Arrows indicate lysed cells. Arrowheads indicate chained cells. (C) Representative cells showing a variety of morphological defects: bent, lysis, chaining with septa, and elongated cells with few septa. Size bar = 10 µm. These results are representative of at least two independent experiments.

## Discussion

In summary, we have built and tested two new tools that we hope will be useful additions to the *C. difficile* molecular biology toolbox. We cloned the regulatory elements of the *C. difficile* xylose utilization operon to generate a xylose-inducible expression plasmid. When driving the expression of *mCherryOpt*, induction is 100-fold or higher and uniform across the population. Bacterial growth was not affected up to 3% xylose, the highest concentration tested. The lack of inducer toxicity is a potential advantage over existing tetracycline- and nisin-inducible vectors (8, 10). In addition, P_*xyl*_ can be combined with tetracycline- and/or nisin-inducible promoters to independently regulate expression of multiple genes in the same cell. Catabolite repression does not appear to be an issue, as the presence of 1% glucose only reduced fluorescence by 40%. A potential caveat is that consumption of xylose might change inducer concentration over the course of an experiment, although this issue could presumably be addressed by deleting the *xylBA* which likely encode products necessary for xylose utilization.

We also constructed a CRISPR interference tool for use in *C. difficile*. The CRISPRi plasmid is an *E. coli*-*C. difficile* shuttle vector that expresses sgRNA constitutively from the P_*gdh*_ promoter and a codon-optimized *dCas9* under control of P_*xyl*_. Repression of gene expression can be tuned by inducing *dCas9* expression with different amounts of xylose. It is relatively easy to target repression to a gene of choice by replacing the 20-nt base-pairing region in the sgRNA with one that is complementary to the gene of interest. Indeed, all of the sgRNAs tested in this study were highly effective (3 targeting *ftsZ*, 2 targeting *slpA*, and 2 targeting *pbp_0712*).

An important limitation of CRISPRi is polarity on downstream (and in some cases upstream) genes (23). In this respect CRISPRi is inferior to in-frame deletions. Deletion and insertion mutants are also preferable for animal studies, where issues of plasmid stability and the requirement for xylose induction make the current CRISPRi tools a poor fit. These limitations might be overcome in the future by integrating constitutively expressed CRISPRi systems into the chromosome.

Despite these limitations, CRISPRi offers several advantages compared to existing methods for constructing null mutants in *C. difficile*. First, it does not require working in a special *C. difficile* background. Second, it is very fast because targeting a new gene is simply a matter of cloning a new sgRNA. A third major advantage of CRISPRi is that it can be used to explore the consequences of inactivating essential genes, as illustrated here for *ftsZ* (depletion phenotype filamentation) and *pbp-0712* (depletion phenotype a complex mixture of lysis and aberrant morphologies). Finally, CRISPRi could be used to poise gene expression at suboptimal levels, with potential applications in drug discovery screens for synthetic phenotypes (23, 44, 45).

## Materials and methods

### Strains, media and growth conditions

Bacterial strains are listed in Table 1. *C. difficile* strains used in this study were either derived from 630Δ*erm* or R20291, both of which have been sequenced. *C. difficile* was routinely grown in Tryptone Yeast (TY) media, supplemented as needed with thiamphenicol at 10 μg/ml (Thi_10_), kanamycin at 50 μg/ml, or cefoxitin at 16 μg/ml. TY consisted of 3% tryptone, 2% yeast extract, and 2% agar (for solid media). *C. difficile* strains were maintained at 37°C in an anaerobic chamber (Coy Laboratory products) in an atmosphere of 10% H_2_, 5% CO_2_, and 85% N_2_. *Escherichia coli* strains were grown in LB medium at 37°C with chloramphenicol at 10 μg/ml or ampicillin at 100 μg/ml as needed. LB contained 1% tryptone, 0.5% yeast extract, 0.5% NaCl and, for plates 1.5% agar.

**TABLE 1.** Strains used in this study

| Strain | Genotype and/or description | Source or reference |
| --- | --- | --- |
| *E. coli* |  |  |
| OmniMAX-2 T1<sup>R</sup> | F' {*proAB*<sup>+</sup> *lacI*<sup>q</sup> *lacZ*ΔM15 Tn10(Tet<sup>R</sup>) Δ(*ccdAB*)} *mcrA* Δ(*mrr-hsdRMS-mcrBC*) φ80(*lacZ*)ΔM15 Δ(*lacZYA-argF*) U169 *endA1 recA1 supE44 thi-1 gyrA96 relA1 tonA panD* | Invitrogen |
| HB101/pRK24 | F– *mcrB mrr hsdS20*(r<sub>B</sub><sup>–</sup> m<sub>B</sub><sup>–</sup>) *recA13 leuB6 ara-14 proA2 lacY1 galK2 xyl-5 mtl-1 rpsL20* | (47) |
| *C. difficile* |  |  |
| R20291 | Wild-type *C. difficile* strain from UK outbreak (ribotype 027) |  |
| 630Δ*erm* | Spontaneous erythromycin-sensitive derivative of strain 630 (ribotype 012) | (53) |
| CRG1496 | 630Δ*erm*Δ*pyrE* | (13) |
| UM275 | 630Δ*erm* with P<sub>veg</sub>::*rfp* downstream of *pyrE* | This study |

### Plasmid and strain construction

All plasmids are listed in Table 2; an expanded version of this table which includes additional information relevant to plasmid assembly is provided in the Supplement (Table S1). Plasmids were constructed by isothermal assembly (46) using reagents from New England Biolabs (Ipswich, MA). Regions of plasmids constructed using PCR were verified by DNA sequencing. The oligonucleotide primers used in this work were synthesized by Integrated DNA Technologies (Coralville, IA) and are listed in Table S2. All plasmids were propagated using OmniMax 2-T1R as the cloning host, transformed into HB101/pRK24 (47), and then introduced into *C. difficile* strains by conjugation (48).

**TABLE 2.**
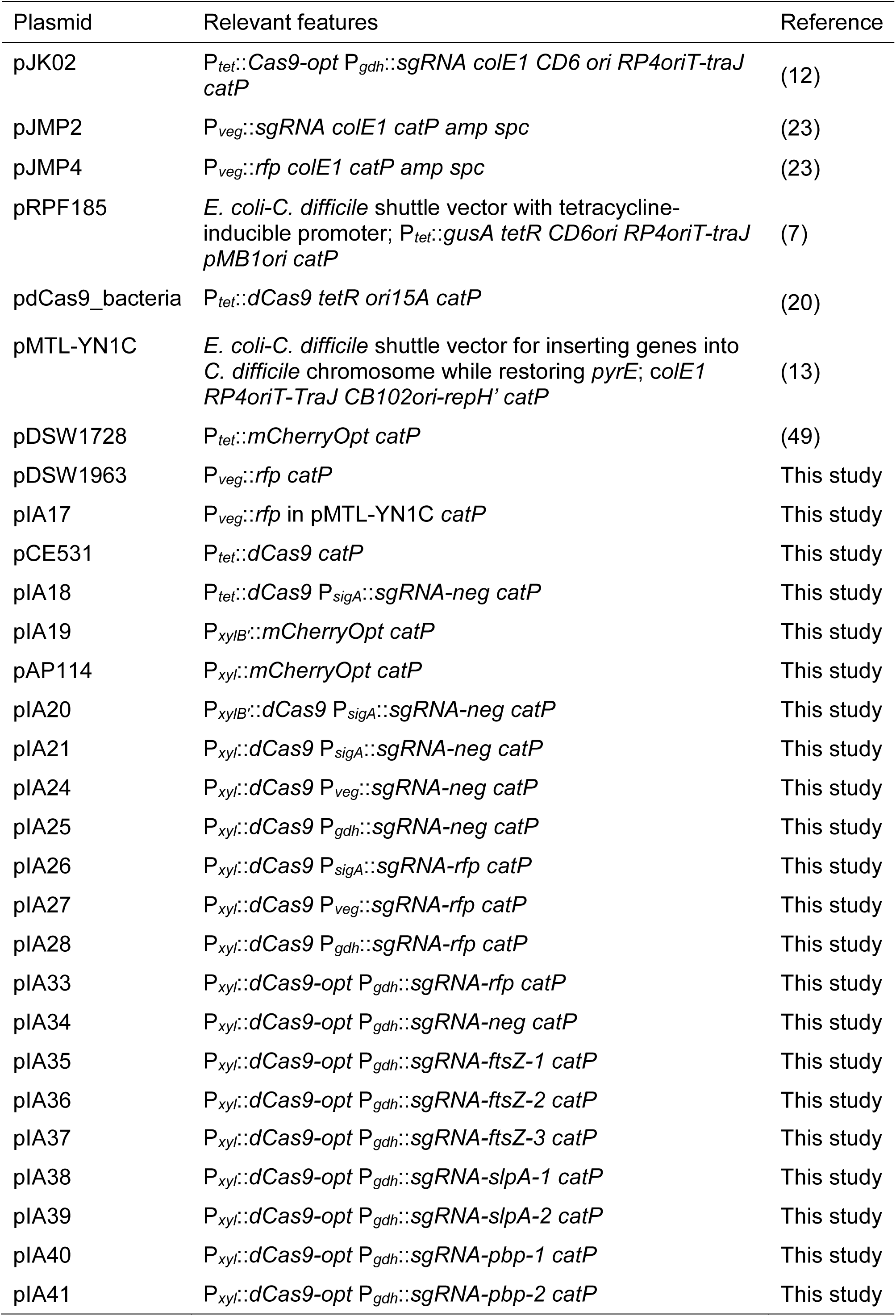
Plasmids used in this study

Plasmid pAP114 is a P_*xyl*_::*mCherryOpt* expression vector derived from pDSW1728 (P_*tet*_::*mCherryOpt*) (49). The *tetR*-P_*tet*_ regulatory element was removed from pDSW1728 by digesting with SacI and BamHI, and replaced with a *xylR*-P_*xyl*_ DNA fragment obtained by PCR of R20291 chromosomal DNA.

CRISPRi plasmids, including the final construct, pIA33, were built on the backbone of pRPF185 (7), an *E. coli-C. difficile* shuttle vector with a chloramphenicol resistance marker and the conjugation locus RP4_*oriT-traJ*_. Initial constructs, harbored *dCas9* from *S. pyogenes* that was amplified from pdCas9_bacteria (20). The final construct, pIA33, harbored *dCas9* with codons optimized for *C. difficile*. This was achieved by amplifying the codon-optimized *Cas9* from pJK02 (12) with primers designed such that the active site residues Asp10 and His840 were changed to alanines. Both versions of *dCas9* were placed under P_*xyl*_ control. Single gRNA (sgRNA) targeting *rfp* was based on the sequence shown to be effective in *B. subtilis* (23) and cloned under control of three different constitutive promoters: a synthetic promoter we termed P_*sigA*_ (28, 50), P_*veg*_ (amplified from pJMP2 (23)) and P_*gdh*_ (amplified from pJK02 (12)), yielding pIA26, pIA27 and pIA28, respectively. The final construct, pIA33, included sgRNA under control of P*_gdh_*. Negative control plasmids (pIA25 for *dCas9* and pIA34 for *dCas9-opt*) had the base-pairing sequence of the sgRNA replaced by 20 nucleotides that do not anneal next to a PAM.

The algorithm provided by Benchling (51) was used to design sgRNAs targeting *ftsZ, slpA,* and *CDR20291_0712*, that last of which codes for a penicillin binding protein, referred to as PBP-0712 (or *pbp-0712)* in this manuscript. Guide parameters were set to default conditions to identify a 20 nucleotide-guide with the PAM set to NGG. Final candidates were selected to be high scoring and bind to the non-coding strand in the first third of the gene sequence. The sequences for sgRNAs are summarized in Table S3.

We constructed a derivative of *C. difficile* 630Δ*erm* that expresses red fluorescent protein (*rfp*) from the P_*veg*_ promoter and is in single copy on the chromosome. This enabled an easy quantitative readout to evaluate our initial, exploratory CRISPRi plasmids. The strain was constructed by allelic exchange (13) using *C. difficile* CRG1496 (630Δ*erm* Δ*pyrE*) as a *pyrE*-deficient recipient and plasmid pIA17 to cross-in P*_veg_*::*rfp* while reconstituting a functional *pyrE*.

#### Xylose induction

Overnight cultures of R20291 or 630Δ*erm* harboring pAP114 were subcultured to an OD_600_ of 0.05 and grown in the presence of varied xylose concentrations at 37°C. Once cultures reached an OD_600_ of about 0.5 (approximately 5 hours), cultures were fixed, and fluorescence quantitated in a plate reader or by flow cytometry. Experiments to evaluate CRISPRi constructs targeting *rfp* followed the same growth protocol unless noted otherwise

#### Fixation protocol

Cells were fixed as previously described (10, 43). Briefly, a 500-μl aliquot of cells in growth medium was added directly to a microcentrifuge tube containing 120 μl of a 5X fixation cocktail: 100 μl of 16% (wt/vol) paraformaldehyde aqueous solution (Alfa Aesar, Ward Hill, MA) and 20 μl of 1 M NaPO_4_ buffer (pH 7.4). The sample was mixed, removed from the Coy chamber, and incubated at room temperature in the dark for 60 min. The fixed cells were washed three times with PBS, resuspended in 110 μl of PBS, and left in the dark for a minimum of 3 hours to allow for chromophore maturation.

#### Microscopy

Microscopy was performed as described previously (10). Cells were immobilized using thin agarose pads (1%). Phase-contrast and fluorescence micrographs were recorded on an Olympus BX60 microscope equipped with a ×100 UPlanApo objective (numerical aperture, 1.35). For FM4-64 red fluorescence the filter set (catalog no. 41004 from Chroma Technology Corp. in Brattleboro, VT) comprised a 538 to 582-nm excitation filter, a 595-nm dichroic mirror (long pass), and a 582 to 682-nm emission filter. Micrographs were captured with a Hamamatsu ORCA Flash 4.0 V2+ CMOS camera.

#### Staining membranes with FM4-64

When membrane morphology was evaluated, cells were stained with the lipophilic dye FM4-64 (Life Technologies). For this, 50 μL of cell culture was removed from the anaerobic chamber and pelleted by centrifugation. After 45 μL of the supernatant was discarded, the pellet was resuspended in the remaining culture fluid and 2 μL of 0.05 mg/mL FM4-64 were added. Cells were then imaged directly with no washing steps.

#### Flow cytometry

Cells were analyzed at the Flow Cytometry Facility at the University of Iowa using the Becton Dickinson LSR II with a 561nm laser, 610/20 bandpass filter, and 600 LP dichroic filter as previously described (10). Data was analyzed using BD FACSDiva Software.

#### Fluorescence measurements with a plate reader

The Infinite M200 Pro plate reader (Tecan, Research Triangle Park, NC) was used to measure bulk samples from cultures as described previously (10). Fixed cells in PBS (100 μl) were added to the well of a 96-well microtiter plate (black, flat optical bottom). Fluorescence was recorded as follows: excitation, 554 nm; emission, 610 nm; gain setting at 160 to evaluate plasmid-based *rfp* expression, or setting 220 to evaluate *rfp* expression from a single, chromosomal copy. The cell density (OD_600_) was also recorded and used to normalize the fluorescence reading.

#### Viability assay

Initial evaluation of CRISPRi targeting *ftsZ, slpA,* and *pbp-0712* was performed by making a serial dilution of an overnight culture and spotting 5 μL on TY Thi_10_ agar with and without 1% xylose. Plates were photographed after overnight incubation (~18h).

#### Evaluation of SlpA levels

S-layer proteins were extracted using low-pH glycine as described (32). Briefly, 40 mL TY Thi_10_ were inoculated 1:50 with an overnight culture and grown for 8h with and without 1% xylose. Cells were removed from the anaerobic chamber, pelleted and washed with 10 mL PBS. Washed pellets were resuspended in 400 μL 0.2 M glycine, pH 2.2, and agitated gently on a tube rotator at room temperature for 30 min. Samples were neutralized with 1 M TrisHCl, pH 8.8, and analyzed by SDS-PAGE using standard methods. Whole cells samples for analysis by SDS-PAGE were prepared by pelleting 1 mL stationary phase culture, resuspending in 80 μL Laemmli buffer, and heating to 95°C for 5 min.

#### MIC determination

Sensitivity to lysozyme was determined by preparing a dilution series of lysozyme (RPI, cat #L38100) in a 96-well plate, in 50 μL TY Thi_10_ with concentrations ranging from 0.5-32 mg/mL. Wells were inoculated with 50 μL of dilute culture suspension (10^6^ CFU/mL, roughly OD_600_ of 0.005) and grown at 37°C for 17h, exposing cells to a lysozyme range of 0.25-16 mg/mL. Unfortunately, high lysozyme concentrations (1 mg/mL or higher) in TY media turns sufficiently turbid to prevent a direct readout of cell growth by optical density. Instead, CFUs were determined as follows. A 10 μL sample from each well was diluted 10-fold by adding to a daughter plate containing 90 μL TY per well. From this, 5 μL were spotted on TY agar and incubated for 20-24 h. The MIC was considered to be the lowest concentration of lysozyme at which 5 or fewer colonies were found.

#### Sporulation assay

Determining sporulation efficiency followed an established protocol (52) with minor modifications. An aliquot (30 μL) of an overnight culture was struck (5 cm) on TY Thi_10_ plates with or without 1% xylose. After 24h growth at 37°C, cells were scraped into 600 μL PBS and vortexed thoroughly. A 300 μL aliquot was removed from the anaerobic chamber, heated to 60°C for 30 min to kill vegetative cells and returned to anaerobic conditions. A 10-fold dilution series in TY was generated from both the untreated and the heat-treated aliquots, followed by spotting 5 μL on to TY agar plates amended with 0.1% taurocholate and 0.1% cysteine to promote germination of spores. Sporulation efficiency was calculated from the viability of the heat-treated samples divided by the viability of the untreated samples.

## Acknowledgments

This work was supported by National Institutes of Health grants R01AI087834 (CDE) and R21AI121576 (CDE and DSW) from the National Institutes for Allergy and Infectious Disease, and by a Development Grant to DSW from the Department of Microbiology and Immunology at The University of Iowa. Some data presented herein were obtained at the Flow Cytometry Facility, which is a Carver College of Medicine / Holden Comprehensive Cancer Center core research facility at the University of Iowa. The facility is funded through user fees and the generous financial support of the Carver College of Medicine, Holden Comprehensive Cancer Center, and Iowa City Veteran's Administration Medical Center. Plasmid pdCas9-bacteria was a gift from Stanley Qi (Addgene plasmid # 44249). We thank Joe Sorg for pJK02, Jason Peters for pJMP02 and pJMP04, Nigel Minton for pMTL-YN1C, and members of the Ellermeier and Weiss laboratories for helpful discussions.

